# Direct RNA sequencing reveals structural differences between transcript isoforms

**DOI:** 10.1101/2020.06.11.147223

**Authors:** Jong Ghut Ashley Aw, Shaun W. Lim, Jia Xu Wang, Yang Shen, Pornchai Kaewsapsak, Eng Piew Louis Kok, Chenhao Li, Boon Hsi Ng, Leah A. Vardy, Meng How Tan, Niranjan Nagarajan, Yue Wan

**Author notes:** These authors contribute equally (Jong Ghut Ashley Aw, Shaun W. Lim, Jia Xu Wang). Correspondence should be addressed to: Y.W. or N.N.

## Abstract

The ability to correctly assign structure information to an individual transcript in a continuous and phased manner is critical to understanding RNA function. RNA structure play important roles in every step of an RNA’s lifecycle, however current short-read high throughput RNA structure mapping strategies are long, complex and cannot assign unique structures to individual gene-linked isoforms in shared sequences. To address these limitations, we present an approach that combines structure probing with SHAPE-like compound NAI-N3, nanopore direct RNA sequencing, and one-class support vector machines to detect secondary structures on near full-length RNAs (*PORE-cupine*). *PORE-cupine* provides rapid, direct, accurate and robust structure information along known RNAs and recapitulates global structural features in human embryonic stem cells. The majority of gene-linked isoforms showed structural differences in shared sequences both local and distal to the alternative splice site, highlighting the importance of long-read sequencing for phasing of structures. Structural differences between gene-linked isoforms are associated with differential translation efficiencies globally, highlighting the role of structure as a pervasive mechanism for regulating isoform-specific gene expression inside cells.

## Introductions

RNAs can fold into complex secondary and tertiary structures to regulate every step of its life cycle^1^. The ability to assign correct structures to the right transcript is key to understanding RNA based gene regulation. Recent progress in RNA secondary structure mapping, using a variety of enzymatic and chemical probes coupled with high-throughput sequencing has resulted in the generation of large-scale RNA structure information across transcriptomes both *in vitro* and *in vivo*^2-10^. This has yielded insights into the pervasive regulatory roles of structure in diverse organisms and cellular conditions^1, 11^. While powerful, current high-throughput structure mapping approaches suffer from drawbacks such as complex library preparation protocols and the lack of structural information for full-length transcripts due to short-read sequencing. As short-reads cannot distinguish structures in shared regions between isoforms, this poses a challenge in our ability to correct assign RNA structural information in individual gene-linked isoforms, limiting our understanding of the role of structure in gene regulation.

Recent developments in long-read, amplification-free sequencing have enabled direct RNA and DNA sequencing by measuring the current across nanopores as single RNA/DNA molecules threads through a biological pore^12^. Additionally, several DNA modifications have been shown to perturb the current across pores differently from typical bases, leading to decodable signal anomalies that reveal both the position and identity of modifications along the genome^13^. In principle, RNA modifications are also decodable but their use for studying full-length RNA structure has yet to be explored^12^.

Here, we couple chemical structure modifications with direct RNA sequencing on nanopores to identify structural patterns in the transcriptome of human embryonic stem cells. Our method, *PORE-cupine* (for *c*hemical *u*tilized *p*robing *i*nterrogated using *n*anopor*es*), identifies single-stranded bases along an RNA by detecting current changes induced by structure modifications (**Figure 1a**). As the method involves only a simple two-step ligation protocol (with a preparation time of two hours prior to sequencing) without the need for PCR-amplification, *PORE-cupine* captures structural information in a transcriptome rapidly and directly. The nature of long-read sequencing through nanopores also allows us to accurately assign and capture structures and their connectivity along individual gene-linked isoforms, deepening our understanding of how the complex and extensively spliced transcriptome could take on different structures to regulate gene expression.

**Figure 1.**
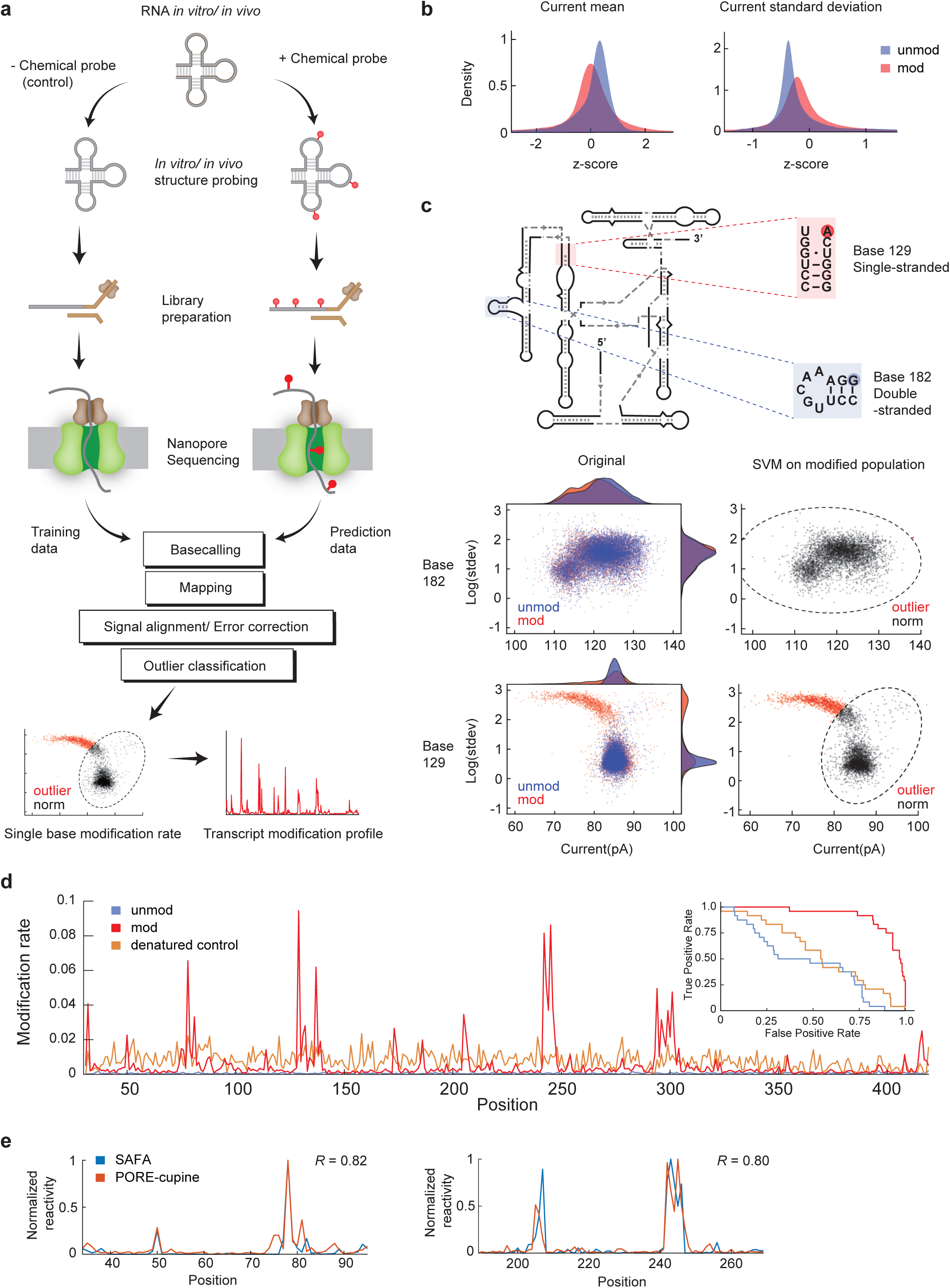
*PORE-cupine* leverages machine learning to profile RNA secondary structure. **a**, Schematic of direct RNA sequencing followed by signal processing. RNAs are structure probed and nanopore sequenced, yielding characteristic voltage signals. The non-structure probed signal is used as a training set to predict modifications from the structure-probed dataset. **b**, Normalized current mean and standard deviation distributions for single-stranded regions on Tetrahymena RNA modified with NAI-N3. With footprinting gels as a guide, the top 10% of the single-stranded regions on Tetrahymena RNA are chosen for these plots. **c**, *Upper*, schematic of the secondary structure of Tetrahymena RNA and the location of a representative single- and double-stranded base. *Lower*, Current mean and log_10_ (standard deviation of current) of the highlighted bases in *upper*. Each datapoint is from a base in a single RNA strand. The leftmost boxes show the distributions for non-structure probed (blue) and structure probed (red) bases before SVM classification. The rightmost boxes show only the structure probed distributions, but with the SVM boundary drawn (dotted lines). Outliers are in red and points within the boundary are in black. **d**, NAI-N3 modification profile along the entire length of the Tetrahymena RNA (red). The modification profile of a randomly modified linearized control (brown) and another non-structure probed replicate (blue) is also shown. The Y-axis indicates the modification rate per base while the X-axis indicates the position along the RNA. Inset: ROC curve for unmodified, modified and denaturing control Tetrahymena sequences. **e**, Comparison of NAI-N3 Modification profile (brown) against SAFA footprinting signal (blue) from two Tetrahymena regions.

## Results

### Structure probing can provide distinct signals detectable using direct RNA sequencing

Numerous chemical probes can penetrate the cellular membrane to modify single-stranded RNA bases inside cells^14^. To determine which of the *in vivo* chemical probes can result in a detectable signal change within nanopores, we tested five different structure probing compounds that modify single-stranded bases (**Supp. Figure 1a**). These include SHAPE and SHAPE-like reagents that acylate the 2’OH of flexible bases: N-methylisatoic anhydride (NMIA), 2-methylnicotinic acid imidazolide (NAI) and 2-methylnicotinic acid imidazolide azide (NAI-N3), as well as base-specific chemical probes: dimethyl sulfate (DMS) and 1-cyclohexyl-3-(2-morpholinoethyl) carbodiimide metho-p-toluenesulfonate (CMCT) (**Supp. Figure 1b**)^6, 14-16^. DMS alkylates single-stranded bases, specifically at N1 of A, N3 of C and N7 of Gs, while CMCT primarily reacts with single-stranded Us at N3 and Gs at N1 positions (**Supp. Figure 1b**). We first performed structure probing using each of these chemicals on the Tetrahymena ribozyme, which has a well-defined secondary structure^17^. Structure probed and non-structure probed Tetrahymena ribozyme RNA were ligated to an adapter and attached to a motor protein before being directly sequenced on the nanopore MinION system^12^ (**Figure 1a**).

As bases with modifications can be miscalled during basecalling, resulting in increased error rates, we tested whether our modified libraries have increased error rates as compared to unmodified libraries. Indeed, we observed a higher base error rate (modified: 6.45%-11.7%, unmodified: 5.26%) in structure modified Tetrahymena sequences for all of the five chemical compounds, with DMS modifications having the highest error rates (**Supp. Figure 2a**). Bases with increased error rates are enriched in single-stranded regions for four out of five chemical modifications, with those modified by NAI-N3 having the highest accuracy rate of around 60 percent (**Supp. Figure 2b**), suggesting that NAI-N3 could be a suitable structure probe to couple with direct RNA sequencing.

**Figure 2.**
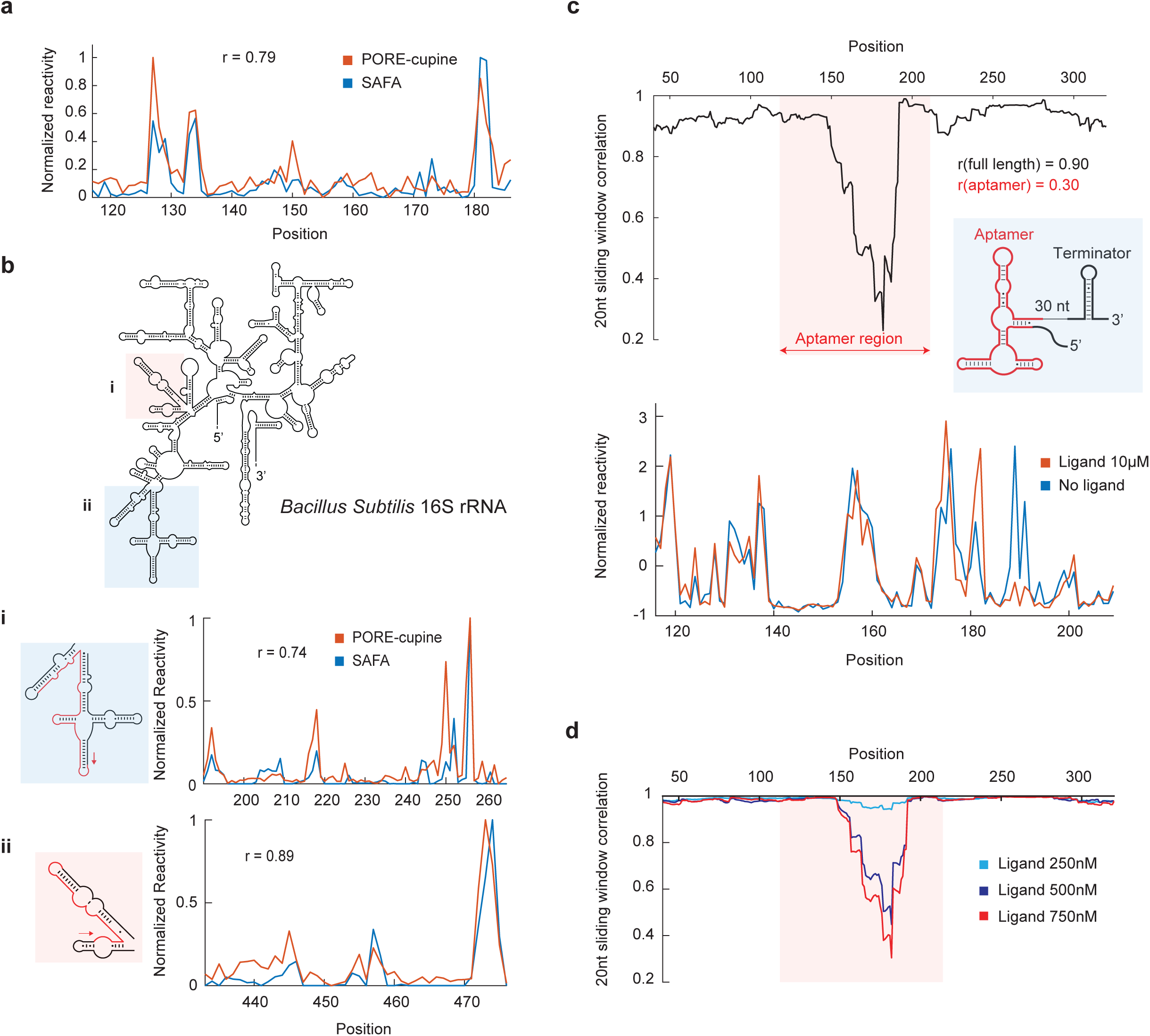
*PORE-cupine* performs accurately *in vitro* and *in vivo*, and captures riboswitch structural dynamics. **a**, Line graph of NAI-N3 modification profile against SAFA footprinting signal from a Lysine riboswitch RNA. The Y-axis indicates normalized reactivity (nanopore) or SAFA signal (footprinting) and the X-axis indicates the position along the RNA. **b**, *Upper*, secondary structure model of 16S rRNA^19^. *Lower*, Line graphs of NAI-N3 modification profile against SAFA footprinting signal from *in vivo* modified 16S rRNA. Red arrow indicates location of RT primer and direction of cDNA synthesis. **c**, *PORE-cupine* captures TPP riboswitch dynamics. 20-nucleotide sliding window Pearson correlation (*upper*) and normalized reactivity profiles of the aptamer region (*lower*) of TPP riboswitch RNA folded in the presence and absence of 10µM TPP. **d**, *PORE-cupine* detects a gradated change in TPP structure. 20-nucleotide sliding window Pearson correlation profiles of TPP riboswitches folded in the presence of increasing concentrations of ligand.

### *PORE-cupine* accurately interprets NAI-N3 signals with SVMs to predict RNA structure

To detect NAI-N3 signals with higher accuracy, we next tested the ability of one class support vector machine (SVM) to perform anomaly detection on our structure probed RNA, based on unmodified sequences. We first utilized the program nanopolish to align the sequenced signal to the RNA sequence (**Supp. Figure 3**)^13^, before extracting two features of the current - the current mean and standard deviation – to determine their distribution in single-stranded versus double-stranded bases in the Tetrahymena RNA. We observed that modified single-stranded bases undergo signal shifts, while double-stranded bases do not, indicating that we can distinguish modification status based on these two features (**Figure 1b)**. Structure signals from two biological replicates are highly correlated, indicating that our data is reproducible (*R*=0.87, **Supp. Figure 4a)**. Using SVM, we are able to identify outliers in single-stranded bases in a folded, NAI-N3 modified, Tetrahymena ribozyme with high accuracy (**Figure 1c-e**). These outliers are not seen in additional biological replicates of unmodified RNA and are evenly distributed in modified denatured RNA in a non-structure specific manner, further indicating that defined peaks in folded modified RNA are due to real structure modifications (**Figure 1d**).

**Figure 3.**
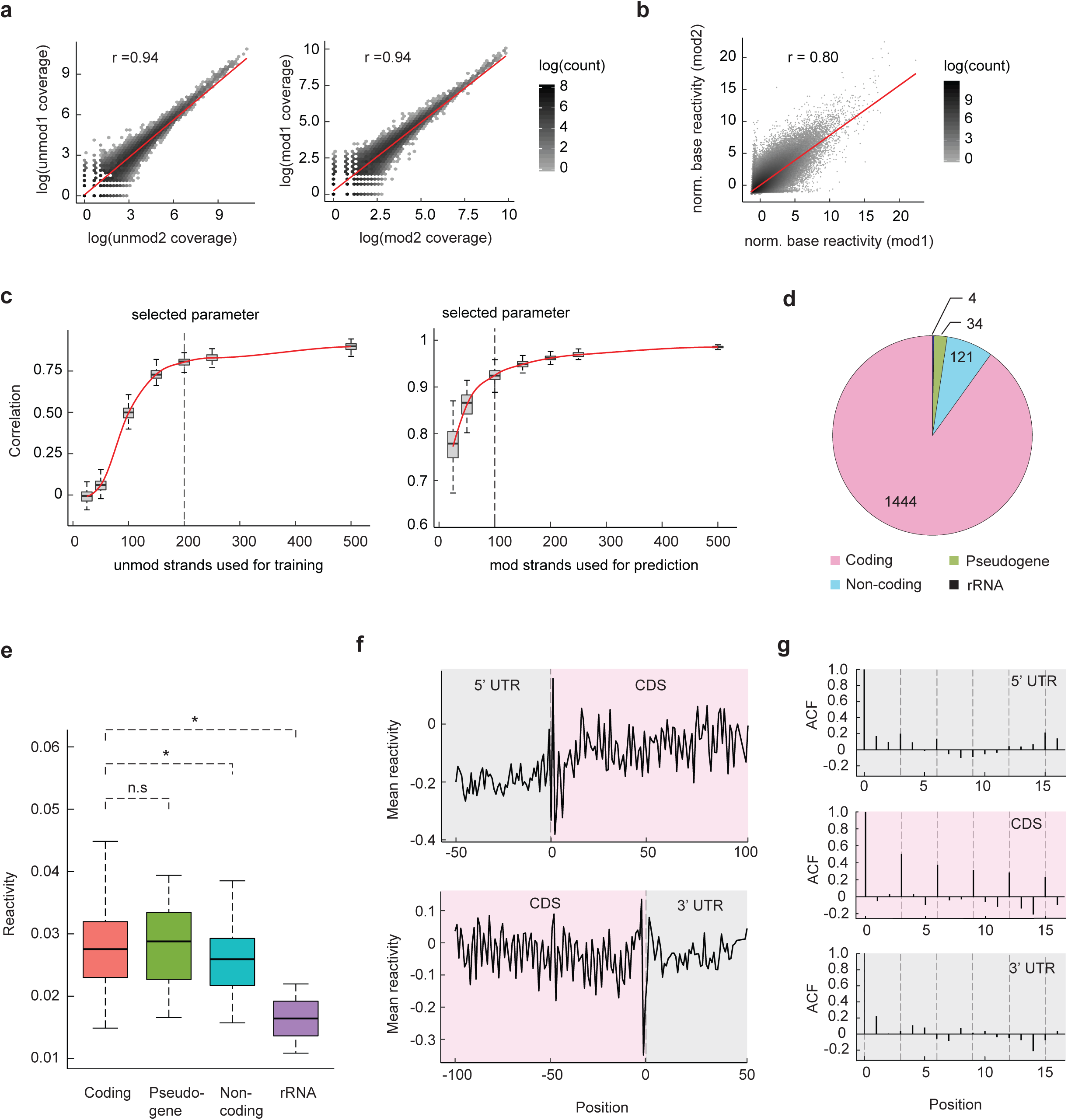
Genome-wide structure analysis in hESC with *PORE-cupine* confirms known global structural features. **a**, Scatterplots showing the numbers of reads/transcript between two biological replicates of unmodified (*left*) and modified (*right*) hESC libraries. **b**, Scatterplot showing normalized base reactivity between two biological replicates of structure probed libraries. **c**, *Left*, Pearson correlation of Tetrahymena RNA structural profiles obtained from subsampling non-structure probed strands (used for training) to the structural profile obtained from the full dataset. The number of structure probed strands used for prediction is 200. *Right*, Pearson correlation of Tetrahymena RNA structural profiles obtained from subsampling structure probed strands (used for prediction) to the structural profile obtained from the full dataset. The number of non-structure probed strands subsampled used for training is 200. **d**, Pie charts showing the distribution of genes belonging to different classes of transcripts after nanopore reads have been processed and filtered. **e**, Median reactivity of different classes of transcripts. P-values are calculated using the two tailed Wilcoxon Rank Sum test with *P*-values of 0.9824, 0.02332 and 0.04389 for the pseudo-gene, non-coding and rRNA classes compared to the coding class. **f**, Metagene analysis of structure aligned according to start (*Upper*) and stop (*Lower*) codons. *PORE-cupine*-derived mean reactivities of selected regions (5’ UTR, 60 nt upstream of the start codon with CDS, 100 nt downstream of the start codon, *upper*, and CDS, 110nt upstream of the stop codon with 3’ UTR, 50nt downstream of the stop codon, *lower*) are shown with these regions. **g**, Metagene autocorrelation function (ACF) plot for the 5’ UTR, CDS and 3’ UTR. Only the CDS contains a periodicity of 3 nucleotides.

**Figure 4.**
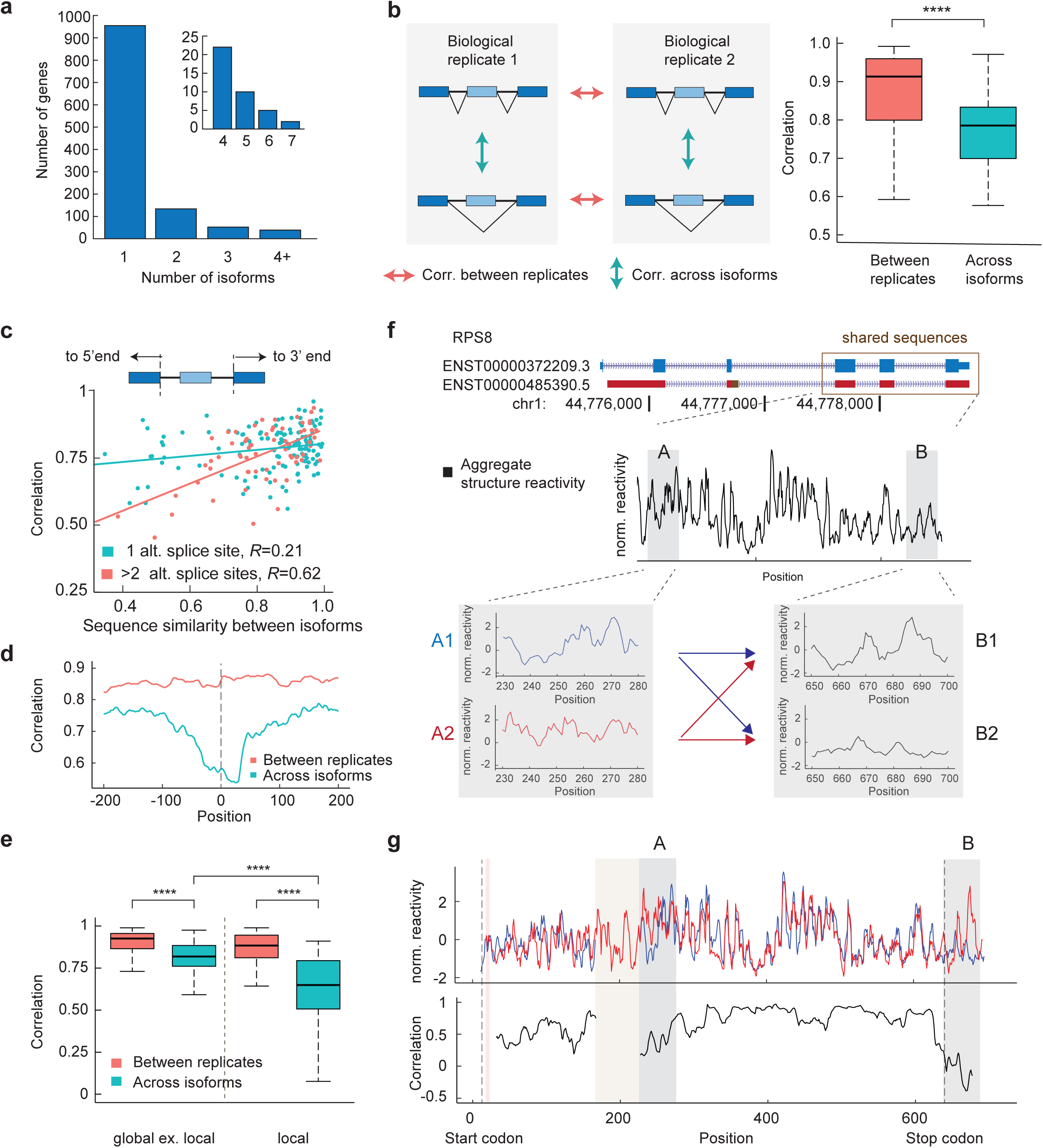
*PORE-cupine* reveals structural differences in shared exons between alternative isoforms. **a**, Histogram showing distribution of structure probed genes according to the number of isoforms present. **b**, *Left*, schematic showing the pair-wise structure comparisons between 1) biological replicates of the same transcript and 2) different isoforms of the same gene. *Right*, boxplot showing distribution of structure similarity for the same isoform across biological replicates (salmon) or between different isoforms within the same biological replicate (teal). *P*-value is calculated using two tailed Wilcoxon Rank Sum test with *P*-values < 2.2e-16. **c**, Scatterplot of structure similarity between shared exons of different isoforms versus their sequence similarity from both sides of the alternative splice site to the 5’ and 3’ end of the transcript. Spearman’s *R* is shown. **d**, Metagene analysis of structural similarity between alternatively spliced isoforms centered at the alternative splice site (blue), as well as structural similarity between biological replicates of the same transcripts centered at the same splice sites (red). Alternative splicing strongly affects RNA structures immediately around the splice site and up to 200 bases and beyond. **e**, Boxplot showing distribution of structure similarity for the same isoform across biological replicates (salmon) or between different isoforms within the same biological replicate (teal), for global excluding local contexts, and for local contexts. Here, global excluding local contexts is defined as regions extending from both sides of the alternative splice site to the 5’ and 3’ end of the transcript, excluding 200bp to the right and left of the alternative splice site, whereas local contexts are defined as 50bp to the left and right of the alternative splice site. The *P*-values are calculated using two tailed Wilcoxon Rank Sum test with *P*-values < 3.97e-11 for the comparison between same isoforms across biological replicates and different isoforms within the same biological replicate for the global excluding local context, *P*-values < 2.2e-16 for the same comparison across the local context, and *p*-value < 4.8e-8 for the comparison for different isoforms within the same biological replicate across the 2 contexts. **f**, Structural information from 2 different isoforms of RPS8. *Upper*, exon and intron organization displayed with their respective Ensemble spliced transcript IDs. An alternative exon seen in our structural data is colored in brown. The three shared exons near the 3’end of the gene is boxed. *Lower*, normalized reactivity profiles for the aggregate signal of the isoforms in the three shared exons. The two structure changing regions (A and B) between the isoforms identified by PORE-cupine are boxed in grey. Without long-read sequencing information, A1 could be linked to B1 or B2, and A1 could be linked to B1 or B2. **g**, Normalized reactivity profiles across the entire length of the different isoforms (top) and the 30 nucleotide sliding window Pearson correlation of the reactivity profiles (bottom). *PORE-cupine* could assign the correct structures to the different isoforms due to long-read sequencing.

Interestingly, over-modification of Tetrahymena RNA resulted in shorter sequences passing through the pore and poor mappability rates (median length of highly modified RNA = 348, lightly modified = 378, unmodified sequences =379 bases, **Supp. Figure 4b**), suggesting that heavily modified strands could be prematurely ejected from pores during sequencing. To perform structure analysis of near full length transcripts, we filtered against transcripts that are shorter than 75% of the full length transcript (**Supp. Figure 4c**). Using SVM, we then estimated the proportion of modified bases for each nucleotide along the RNA as a “reactivity profile”. *PORE-cupine*’s “reactivity profile” correlates well with traditional RNA footprinting data, indicating that *PORE-cupine* signals are accurate and quantitative (**Figure 1e**).

To further test the accuracy of *PORE-cupine* in other *in vitro* and *in vivo* structure probed RNAs, we sequenced two biological replicates of the lysine riboswitch and 16S rRNA^18, 19^. *PORE-cupine* reactivity profiles were found to be highly correlated between biological replicates (**Supp. Figure 4d**,**e**), and are well correlated with footprinting data, demonstrating that we can detect structures across various RNAs accurately (**Figure 2a**,**b**). We next tested if localized structural changes observed in two RNA confirmations of a riboswitch (with and without ligand) can be detected using *PORE-cupine*^20^. *PORE-cupine* reactivity profiles with and without ligand (10µM TPP) showed high correlation throughout the RNA (*R*=0.9), except for the structure changing aptamer region (*R*=0.3, **Figure 2c**). In addition, we also observed a gradated change in riboswitch structure under lower concentrations of TPP, indicating that *PORE-cupine* signals are specific and semi-quantitative (**Figure 2d**).

### Genome-wide analysis of RNA structures in hESC using *PORE-cupine*

Beyond studying individual transcript structures, groups of transcripts can share similar structures to perform functions inside the cell. As such, studying how RNA structures are regulated inside cells, in a genome-wide manner, is key to understanding gene regulation. We applied *PORE-cupine* to study the RNA structural landscape in human embryonic stem cells (hESC) by sequencing four biological replicates of NAI-N3 structure probed and unprobed hESC transcriptomes totaling to 10 million sequenced reads for each condition (**Supp. Table 2, Supp. Figure 5a**). The median length of the mapped reads are 830 and 902 bases for modified and unmodified strands respectively, covering >60% of an average transcript (**Supp. Figure 5b**). The transcript abundance and reactivity profiles of the RNAs were highly correlated between biological replicates (**Figure 3a,b**), and between modified and unmodified samples, indicating that our data is reproducible and that we capture similar transcript profiles regardless of modification status (**Supp. Figure 5c**).

**Figure 5.**
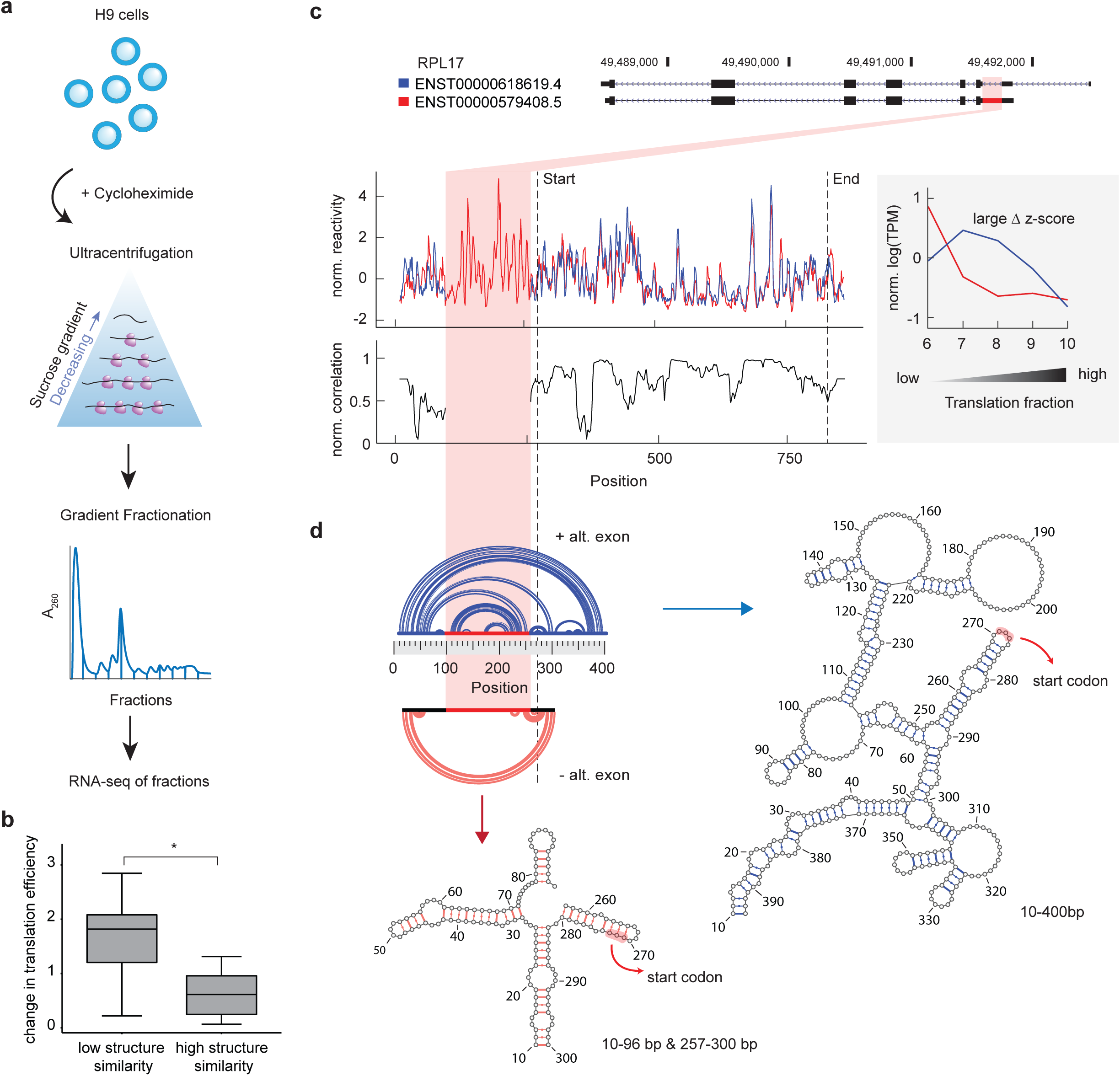
Structure differences between isoforms are correlated with translation efficiency. **a**, Schematic of the TrIP-seq experiment in hESCs^24^. **b**, Boxplot showing that gene-linked isoforms with greater structural similarity show smaller differences in translation efficiency. P-value calculated by two tailed Wilcoxon Ranked Sum Test, *P*-values =0.0482, N_low_=18 and N_high_=5. **c**, Structural information from a pair of isoforms from RPL17 that show translational differences. *Upper*, exon and intron organization displayed with their respective Ensemble spliced transcript IDs. The alternative exon seen in our structural data is highlighted in red. *Lower*, normalized reactivity profiles for the different isoforms and the 30 nucleotide sliding window Pearson correlation of the reactivity profiles. The normalized log(TPM) expression level for the isoforms in fractions 6-10 is shown in the inset. **d**, RNAcofold-derived structures for the region around the alternative exon based on SPLASH read counts. The start codon is highlighted (red) on the structures. The upper structure (blue) is based on RNAcofold-calculated interactions for 10-400bp, while the lower structure (red) is based on the RNAcofold-calculated interactions between regions 10-96bp and 257-300bp.

To determine the number of full-length transcript reads needed for accurate structure determination, we subsampled the number of unmodified and modified Tetrahymena reads used and compared the structural information obtained from the subsampled set to that of the full dataset. As expected, the correlation increases with the number of reads used and begins plateauing around *R*=0.8 at 200 strands of unmodified and 100 strands of modified RNA (**Figure 3c**). At this threshold, we obtained structural information for 1444 coding genes, 121 noncoding genes, 34 pseudogenes and rRNAs across the hESC transcriptome (**Figure 3d**).

To examine whether different classes of RNAs have divergent structural propensities, we averaged the structural signal for each class and observed that rRNAs are the most highly structured, followed by lncRNAs and mRNAs, in agreement with the importance of structure for noncoding RNAs^3^ (**Figure 3e**). Metagene analysis of mRNAs aligned by their translational start and stop sites showed the classic three nucleotide structure periodicity in their coding sequences (CDS), and not in their 5’ and 3’UTRs^2, 10, 21^ (**Figure 3f, g**), highlighting *PORE-cupine*’s ability to recapitulate known patterns in hESC data.

### Detecting structural differences in shared exons from alternative isoforms

The human transcriptome is extensively spliced^22^. This high complexity of the transcriptome makes assigning RNA structural information to individual isoforms extremely challenging as gene-linked isoforms share extensive sequence similarities. As the majority of the short reads fall in shared exons, and cannot be precisely mapped to a specific isoform, short-read sequencing limits our understanding of how isoforms could take on different structures for gene regulation. As recent global RNA structure studies have shown that even small differences between isoforms in the 3’UTR can cause large structural differences with functional consequences in gene regulation^23^, we hypothesize that alternative splicing across the rest of the transcript could also have large impacts on RNA structure in shared regions. To analyze RNA structures present in these isoforms, we mapped our direct RNA sequencing data to transcripts present in the Ensemble database (**Methods**). 101 genes (219 isoform pairs) had at least two or more isoforms that pass our threshold of having at least 200 strands for accurate structure analysis (**Figure 4a, Supp. Figure 6a**) and we performed our downstream analysis on these transcripts.

We observed that the majority (209 out of 219, 95%) of the gene-linked isoforms show structural differences in shared regions. Globally, this is reflected by their lower structural similarities in shared sequences across gene-linked isoforms, as compared to biological replicates of the same transcript (**Figure 4b**). In general, sets of isoforms with greater sequence differences show larger structural differences in shared regions (**Figure 4c**). This correlation is stronger when there are two or more alternative splice sites along a transcript, resulting in wide-spread structure changes across the entire transcript (**Figure 4c**). While the biggest structural differences appear to occur immediately around the alternative splice site, 43% of isoforms contain both local and distal structural changes away from the alternative splice sites (**Figure 4d, Supp. Figure 6b**). We confirmed this distal impact on structures by plotting structure differences in identical sequences that are more than 200 bases away from the alternative splice site (**Figure 4e**).

### *PORE-cupine* can phase structures along long isoforms

The presence of two or more alternative structures that reside and span identical sequences makes it particularly challenging for short-read sequencing to determine which combinations of RNAs structures co-exist in a single isoform. An example of this is RPS8, which is alternatively spliced into two isoforms. The two isoforms share identical sequences for three of the exons towards the end of the sequence, but are alternatively spliced near the 5’ end of the gene (**Figure 4f**). *PORE-cupine* analysis shows that the two isoforms contain different structures (A1 versus A2, and B1 versus B2) that span ∼400 bases from each other in the shared sequences. In the absence of long-read sequencing, the lack of connectivity between structures A and B makes it difficult to know whether A1 is linked to B1 or B2 in the blue isoform and vice versa. However, with long-read sequencing, we are able to link and correctly assign structural information to their individual isoforms in shared regions (**Figure 4g**). Globally, the median distance between any two changing structural regions along an isoform is 268 bases, with 25% of transcripts having two structural changing regions more than 648 bases apart (**Supp. Figure 6c**). This large fraction of transcripts having structures located far apart demonstrates the importance of *PORE-cupine* in phasing structures along long isoforms, providing connectivity in RNA structure information across the transcriptome.

### Isoforms with structural differences show differences in translation efficiency

RNAs can utilize different structures to regulate gene expression, including translation, splicing and decay^1^. To determine the extent to which structural differences between isoforms could regulate differential gene expression, we performed TrIP-seq on hESCs and correlated RNA structural changes between the isoforms with their translational efficiencies (**Figure 5a, Supp. Figure 7a**)^24^. We obtained a high degree of correlation between two biological replicates across polysome fractions (**Supp. Figure 7b**), and observed that highly translated transcripts are found in high ribosomal load fractions, while poorly translated transcripts are found in low ribosomal fractions as expected (**Supp. Figure 7c**), indicating that our TrIP-seq data is robust and provides an accurate reflection of mRNA-polysome association.

We observed that 16% of our structural changing gene-linked isoforms show changes in translation efficiency by TrIP-seq. In particular, alternative splicing in 5’UTRs that result in large structural changes are associated with big translation efficiencies differences between gene-linked isoforms, confirming previous observations that 5’UTR structures are important for regulating translation (**Figure 5b**). One of the genes that we detected in our dataset, RPL17, consists of 58 isoforms in the ensemble database, seven of which are detected in our nanopore dataset. One of the transcripts (ENST00000579408.5) is poorly translated, and contains a retained intron of 161 bases in the 5’UTR (**Figure 5c**, *upper*). To study how this retained intron results in structural changes in the 5’UTR and in translation repression, we examined pair-wise RNA interactions within this region identified by SPLASH^25^. Interestingly, SPLASH reads showed a very strong interaction between the retained intron with both an upstream and downstream sequence, resulting in an extensively paired structure around the start codon (**Figure 5d**), sequestering it in a highly structured environment. In the absence of the retained intron, the isoforms fold into a simpler structure, allowing the start codon to be more accessible for translation.

## Discussion

The human transcriptome is both complex and tightly regulated by numerous sequence and structural features along each transcript. Assigning correct structural information to individual transcripts is the first step in understanding structure-based gene regulation. Here, we coupled RNA structure probing with direct RNA sequencing on nanopores to better understand structure-function relationships in the cell. Modifications on single-stranded bases can be detected using SVMs and the frequency of outliers at each base can be measured in terms of reactivity at the base to serve as a proxy for single-strandedness along an RNA. Compared to short-read high throughput sequencing protocols, which are typically time consuming^2, 6, 26, 27^, *PORE-cupine* provides a fast and direct way to assay for RNA structures and dynamics genome-wide. As nanopore sequencing allows long strands of RNAs to be sequenced, *PORE-cupine* is also able to obtain structure information along large stretches of RNAs, enabling us to determine and phase RNA structures in different isoforms. Interestingly, many isoforms show structural differences that are correlated with their translational status, strengthening the importance of structure in regulating mRNA translation. *PORE-cupine* expands our current repertoire of RNA structure probing strategies to deepen our understanding of the role of RNA structure in isoform-specific gene regulation^2, 6, 9, 10, 28^.

## Methods

### RNA modifying reagents

CMCT, NAI, NMIA and DMS were purchased from Sigma Aldrich. NAI-N3 was synthesized as previously described from ethyl 2-methylnicotinate in 4 steps, as in Spitale et al^6^.

### In vitro transcription, folding and *in vitro* structure probing

RNA was transcribed from PCR-amplified inserts using the Hiscribe T7 high yield synthesis kit (NEB). 0.1µg/µl of the RNA of interest was heated to 90°Cfor 2 mins and cooled at 4°Cfor 2 mins before adding chilled 10X RNA structure buffer (500mM Tris pH 7.4, 1.5M NaCl, 100mM MgCl_2_), and TPP ligand if necessary. The RNA was then subject to a slow ramp-up in temperature to 37°Cover 30 mins, after which the RNA-modifying chemical was added and incubated with the RNA for 5 mins (“1x” condition as described in the text—”5x” refers to a 25 mins incubation). Depending on the solvent for the RNA-modifying chemical, DMSO or water was added to the negative control. CMCT, NAI and NAI-N3 were added to final concentrations of 100mM, while NMIA and DMS were used at final concentrations of 20mM and 5%(v/v) respectively. DMS reactions were quenched with 30% β-mercaptoethanol in 0.3M sodium acetate. Reactions were column-purified (Zymo research) and resuspended in nuclease-free water.

### RNA footprint analysis

To determine sites of modifications along an RNA, an RT primer (IDTDNA) was designed around 20bp downstream of the region of interest. Primers were radiolabeled with P^32^, using T4 polynucleotide kinase (NEB) and γ-ATP (PerkinElmer). The reaction product was run out and purified from a 15% TBE-Urea PAGE gel. 1µl of the purified, labelled primer was then incubated with 500ng of RNA (in 5.5µl) at 65°Cfor 5 mins, followed by 35°Cfor 5 mins, and cooled to 4°C. 3µl of SHAPE buffer (1:1:4 ratio of DTT: dNTPs: First-strand buffer) was added together with 0.5µl of Superscript III before bringing the mix to 52°Cfor 10 mins. 0.5µl of NaOH was added to denature the RNA before adding 10µl of RNA loading dye (47.5% formamide, 0.01% SDS, 0.01% bromophenol blue, 0.005% Xylene Cyanol, 0.5mM EDTA) and running on a 8% TBE-Urea PAGE sequencing gel. Gels were dried for 2 hours on a vacuum gel drier before exposure to a phosphorimager plate for 24 hours. The phosphorimager plates were imaged on the Amersham Typhoon 5 Biomolecular Imager (General Electric). Gel images were quantified using the software Semi-automated footprinting analysis (SAFA)^29^.

### Human and bacterial cell culture with *in vivo* SHAPE modification

hESC (H9) cells were obtained from neighbouring labs in the Genome Institute of Singapore. H9 cells were cultivated in feeder-free conditions with mTESR basal media supplemented with mTESR supplement (Stemcell Technologies) and on 15cm Matrigel (BD)-coated tissue culture plates. Cells were seeded at a 1:6 ratio and routinely passaged with Dispase (Stemcell Technologies).

For *in vivo* SHAPE modification, cells were rinsed once on the plate with room temperature PBS (Thermo Fisher Scientific) before being incubated with Accutase (Stemcell Technologies) for 10 mins at 37°Cto dissociate cells. The total volume was then made to 50ml with PBS and then pelleted, after which the cells were washed once more in PBS. The cells were spun down once more at 400g for 5 min before resuspension in 950µl of PBS in a 1.7ml Eppendorf. 50µl of 2M NAI-N3 in DMSO(+) or DMSO(-) only was then added to two separate cell suspensions and immediately mixed by inversion, before incubation at 37°Cat 10rpm (Model 400 hybridization incubator, Scigene) for 5 mins. After the incubation period, the cell suspensions were immediately spun down at 400g for 5 mins at 4°C. The supernatant was removed before total RNA was isolated with the addition of TRIzol reagent (Thermo Fisher Scientific).

*Bacillus Subtilis* strain 168 was grown in LB media at 37°C to an OD_600_ = 0.6. Cells were harvested by centrifugation at 3000rpm for 5 mins, pelleted and washed in PBS, before being treated with 100µM NAI-N3 as in the H9 example above for 5 mins at 37°C(“1x” condition as described in the text—”5x” refers to a 25 minute incubation). The cells were pelleted after incubation and then resuspended in bacteria lysis buffer (1% SDS, 8mM EDTA, 100mM NaCl) and lysozyme (final concentration: 15mg/ml), incubating for 15 mins at 37°C. Total RNA was then isolated with the addition of TRIzol reagent.

### Direct RNA sequencing library preparation

Direct RNA sequencing libraries were constructed using an input of 500ng of RNA for single templates or 1µg of mRNA isolated from H9 hESC total RNA. H9 mRNA was isolated using the Poly(A) purist MAG kit (Thermo Fisher Scientific). Oxford Nanopore Technologies’ direct RNA library preparation kit SQK-RNA001 was used for all sequencing runs in this study. All preparation steps were followed according to the manufacturer’s specifications, except for the omission of a singular reverse transcriptase step. The libraries were loaded onto R9.4.1 or R9.5 flow cells and sequenced on a MinION device. For the in vitro and 16S templates, the RTA DNA adapter in the first step was replaced with a DNA adapter complementary to the 3’ end of the RNA. The sequences of these adapters are detailed in Supplementary Table 1. The sequencing parameters were modified for all runs, and the exact changes can be found in the github link found under software and scripts below.

### Polysome fractionation

H9 cells from a 15cm plate were treated for 10 mins with 100µg/ml cycloheximide at 37°C. The cells were next washed with warm PBS, dissociated with trypsin, and neutralized with ice cold media containing FBS (all supplemented with 100µg/ml cycloheximide). Next, they were pelleted before resuspending in 1x RSB buffer (10mM Tris-HCl pH 7.4, 150mM NaCl, 15mM MgCl_2_) with 200µg/ml cycloheximide, lysed in lysis buffer (10mM Tris-HCl pH 7.4, 150mM NaCl, 15mM MgCl_2_, 1% Triton-X, 2% Tween-20, 1% Deoxycholate), and incubated on ice for 10 mins. Centrifugation at 12,000*g* for 3 mins removed the nuclei, and the supernatant was removed for a subsequent centrifugation.

Equal OD units were loaded onto a linear 10%-50% sucrose gradient that was made using the 107 Gradient Master Ip (BioComp Instruments). The gradients containing the cell lysate were centrifuged in SW41 bucket rotors (Beckman Coulter) at 36,000 rpm at 8°Cfor 2 hours. 12 fractions were separated and collected from the top of the gradient using the PGF Ip piston gradient fractionator (BioComp Instruments) and the Fraction Collector (FC-203B, Gilson). The absorbance readings were collected at 260nm with a Econo UV Monitor (EM-1 220V, Biorad). After fractionation, 110µl of 20% SDS and 12µl of Proteinase K (Thermo Fisher Scientific) were added to each fraction for a 30 min incubation at 42°C, after which GeneChip eukaryotic poly-a RNA controls (Affymetrix) were added to each fraction. Total RNA from each fraction was extracted using phenol-chloroform-isoamyl alcohol (25:24:1, Sigma).

### TrIP-seq library preparation

1µg of total RNA from the polysome fractionated H9 cells was poly-a selected using the Poly(A) mRNA Magnetic Isolation Module (NEB). The mRNA was then adapter ligated and prepared for Illumina sequencing using the Ultra Directional RNA Library Prep Kit (NEB).

### Basecalling and Mapping

Reads were basecalled with Albacore version 2.3.3 without filtering. Basecalled sequences were aligned with Graphmap version 0.5.2^30^. For single gene mapping, references for the individual genes were used. For H9 transcriptome mapping, cDNA and non-coding reference sequences obtained from Ensemble were used (GRCh38).

### Calculation of error rates

Mismapping rates per strand were calculated from aligned bam files with python. Mismapping rates for each modification were analyzed in R and plotted using the ggpubr package. bam-readcount was used to calculate the mismapping rate per position. The fold-differences of error rate per positions were calculated by dividing the error rate of modified samples with that of the unmodified sample. Accuracy of the top 10% of the fold-difference from each samples were used to calculate accuracy by comparison to the secondary structure of the Tetrahymena ribozyme^17^.

### Alignment of current signal

Current measurements above 200pA and below 0pA were deemed as noise and removed from the raw nanopore sequencing file. Subsequently, current signals was aligned with Nanopolish version 0.10.2, and multiple events from the same reads and positions were merged into a single event, by calculating the weighted average of event mean, weighted event standard deviation and sum of event length.

### Calculation of reactivity

One-class support vector machine method was used to calculate the percentage of modification per position. All replicates of unmodified libraries were combined and used for training. Models for individual positions of each transcript were generated with the samples from the unmodified samples as training data. Two features, the event mean and event standard deviation, were selected as features to train and predict perturbations in the signal. Positions with unmodified strands less than 200 and modified strands less than 10 percent of the total coverage were removed. Reactivity scores were calculated by taking the percentage of modifications detected per position, and further z-score normalized.

### H9 transcriptome analysis

Two features, event mean and event standard deviation, were used to train the models for each position of transcripts. The models were then used to detect modification from the modified libraries. Only transcripts that were present in both replicates of the modified libraries, with a minimum coverage of above 100, less than 20% mismatch rates, and transcript length more than 300 nucleotides, were retained for further analysis. A 5-nucleotide moving average was applied to transcript reactivity and then z-score normalized.

### Isoform comparison

Transcripts were aligned to the genomic positions to allow the ease of comparison of reactivity between isoforms. Within an isoform pair, reactivity from the shared exons are kept. For global analysis, all shared positions were used to calculate Pearson correlation. For local analysis, 100 nucleotides left and right of the differential splice sites were used to calculate Pearson correlation. Results with less than 100 nucleotides to the left and right were discarded.

### Software and scripts

Aligned signals were analyzed with R version 3.4.1 and the source code for all scripts can be found at https://github.com/awjga/Porecupine.

## Data availability

## Acknowledgements

We thank members of the Wan lab, Tan lab, Fei Yao, Hwee Meng Loh, Chiea Chuen Khor and Mile Sikic for helpful discussions. YW is supported by funding from A*STAR, Society in Science - Branco Weiss Fellowship, EMBO Young Investigatorship and CIFAR global scholarship.

## Author contributions

Y. Wan conceived the project. Y. Wan, N. Nagarajan, M.H Tan, B.S Ng, L. Vardy designed the experiments and analysis. S. W Lim and J.X Wang performed the experiments with help from P. Kaewsapsak. J.G Aw and Y. Shen performed the computational analysis with help from C. Li and EP. Kok. Y. Wan organized and wrote the paper with J.G Aw and S. W Lim and all other authors.

## Supplementary figure legends

**Supplementary Figure 1. Chemical structures of RNA structure probing compounds and associated reaction products. a**, Chemical structures of RNA structure probing compounds. Side chains for the carbodiimide of CMCT are highlighted and abbreviated as R’ and R’’ for part (**b**). **b**, RNA nucleotide triphosphates with chemical adducts formed from reaction with structure probing compounds. Adducts are highlighted in green.

**Supplementary Figure 2. Statistics of mismapping rates to detect structure modifications on RNA. a**, Boxplots showing the distribution of mismapping rates for different structure probing chemicals. Mismapping rates (not considering indels) for *in vitro* structure probed Tetrahymena RNA are compared against the non-structured probed control. P-values are calculated using the two tailed Wilcoxon Rank Sum test, *P*-values < 2.2e-16, N_unmod_=15154, N_NAI_=6962, N_NMIA_=28427,N_CMCT_=22979,N_NAI-N3_=42107 and N_DMS_=13679. **b**, Barcharts showing the accuracy of mismapping rate for the different compounds. The proportion of the top 25% of mismapping signal is compared against single-stranded bases from the secondary structure map of Tetrahymena RNA. The accuracy rate from a random prediction is shown as a dotted line.

**Supplementary Figure 3. Schematic of the bioinformatic workflow of *PORE-cupine* to obtain reactivity score per base**. Sequenced strands were basecalled using Albacore and mapped to the respective sequences using Graphmap. We used Nanopolish to align the raw signals and to extract current features which were used to train unmodified data using SVM.

**Supplementary Figure 4. *PORE-cupine* sequencing statistics for different RNAs. a**, Scatterplot showing the distribution of normalized base reactivity between two structure probed Tetrahymena RNA samples. **b**, Mapping rates for structure probed versus non-structure probed Tetrahymena RNA. The “5x” RNA sample has been reacted with the same concentration of NAI-N3 for 5 times as long as the “1x” RNA sample, as described in Methods. **c**, Effect of length filters on structure probed and non-structure probed Tetrahymena RNA sequencing data for both the 1X RNA sample (*Upper*) and 5X RNA sample (*Lower*). **d**,**e**, Scatterplot showing the normalized base reactivity between two structure probed 16S rRNA (**d**) and lysine riboswitch (**e**) samples.

**Supplementary Figure 5. Statistics of H9 hESC mRNA *PORE-cupine* sequencing output. a**, Barplots showing the number of total H9 nanopore reads left after basecalling and mapping in treated and untreated human hESC samples. **b**, Boxplots showing the distribution of mapped lengths of untreated (left) and NAI-N3 treated (middle) hESC mRNA. We also added the distribution of expected lengths based on ensemble annotation for comparison (right). **c**, Scatterplot showing the distribution of total reads per transcript for all processed structure probed and non-structure probed transcripts in the hESC transcriptome. *Top left*, non-structure probed (replicate 1) versus non-structure probed (replicate 2); *Top right*, structure probed (replicate 1) versus non-structure probed (replicate 1); *bottom left*, structure probed (replicate 1) versus non-structure probed (replicate 2); *bottom right*, structure probed samples against each other.

**Supplementary Figure 6. RNA structures in different isoforms of the same gene. a**, Structural information from 4 different isoforms of RPLP0. *Upper*, exon and intron organization displayed with their respective Ensemble spliced transcript IDs. Alternative exons seen in our structural data are highlighted in red. *Lower*, normalized reactivity profiles for the different isoforms and their aggregate signal. **b**, Structural information from RACK1. *Upper*, exon and intron organization displayed with their respective Ensemble spliced transcript IDs. Alternative exon is seen in red (also in inset). Lower, normalized reactivity profiles for 2 different isoforms of RACK1, their aggregate signal, and the 30 nucleotide sliding window Pearson correlation between the isoforms. Two regions of low correlation, one at the alternative splice site, and another around the 550 to 570 nucleotide window are shown. **c**, density plot showing the distribution of distances between any two changing regions along an isoform. The median distance is 268 bases, demonstrating the need for long-read sequencing to connect the structure changes.

**Supplementary Figure 7. TrIP-seq data is robust and provides an accurate reflection of mRNA-polysome association. a**, Gradient profile of polysome fractions collected. **b**, Pair-wise correlations of the read-counts/transcript for each fraction between 2 biological replicates. Fractions and batches are denoted as F2-12 and B1-2 respectively. **c**, Distribution of read-counts across different polysome fractions for a highly translated RNA (Actin B, *left*) and a poorly translated RNA (Activating transcription factor 4, *right*).

