## Supplementary figures and images for "Direct RNA sequencing reveals structural differences between transcript isoforms"

### Supplemental Figures

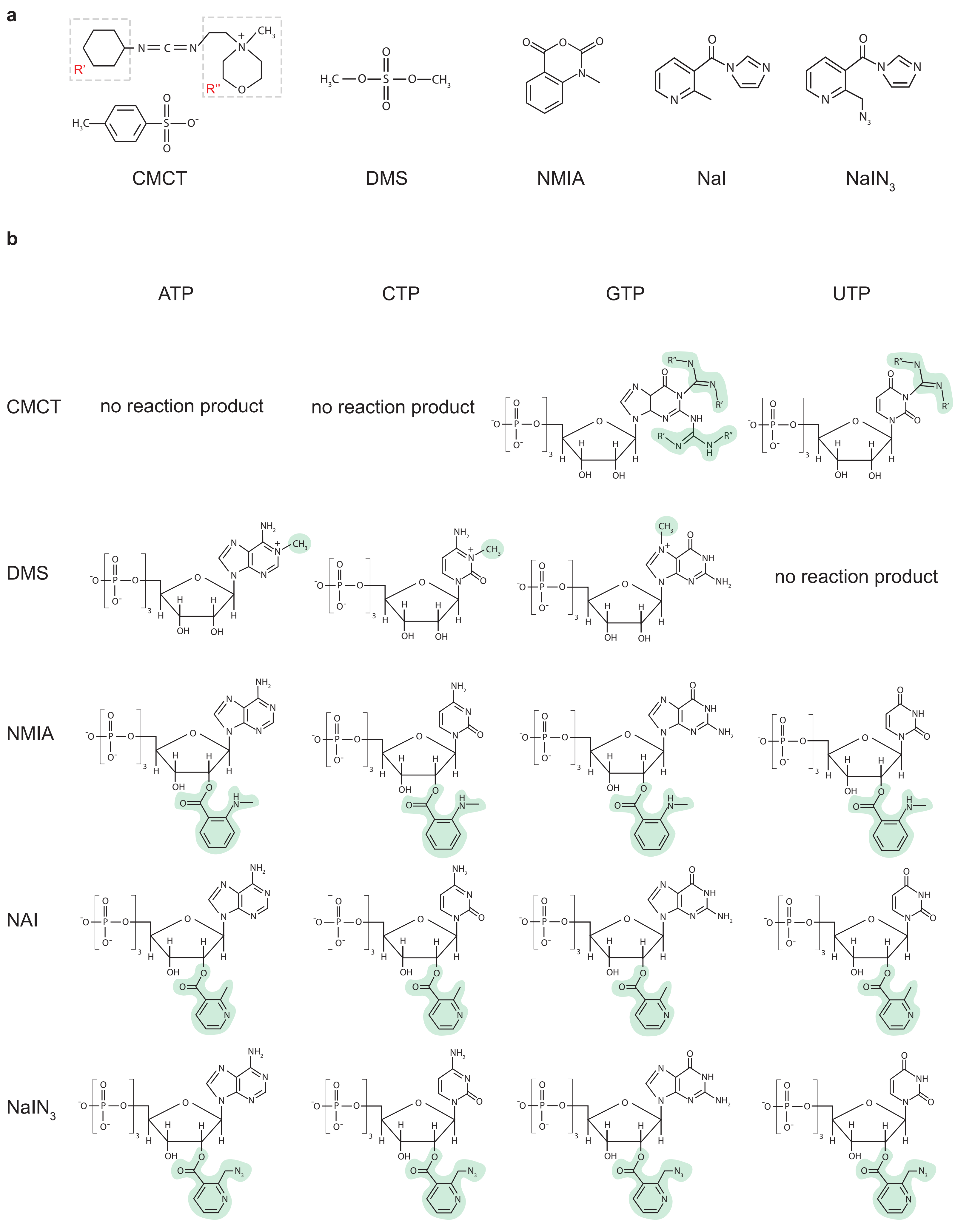

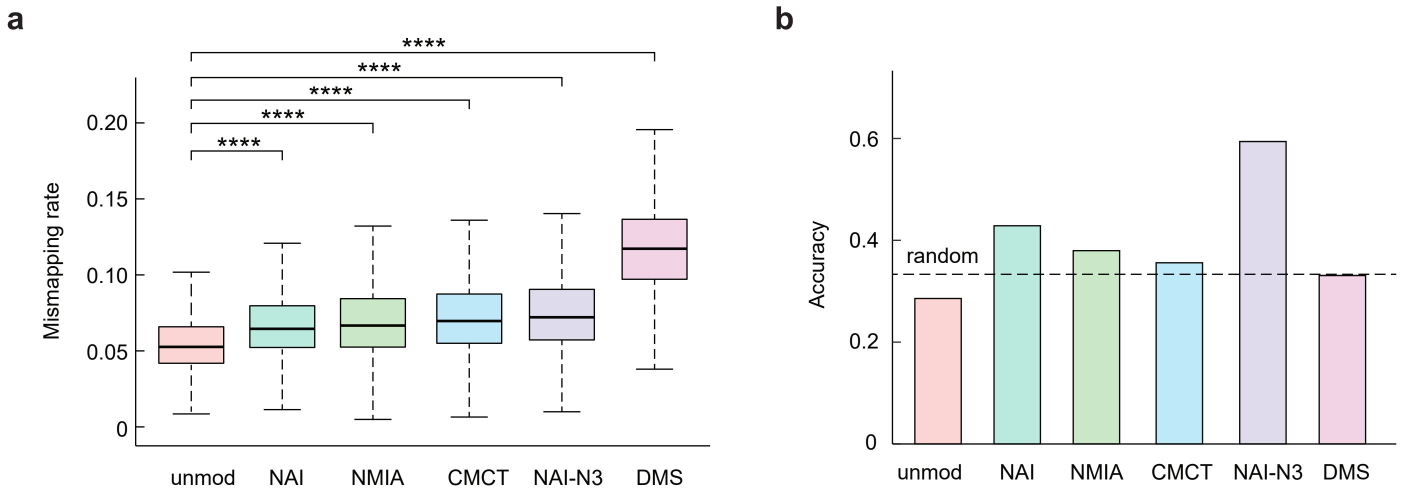

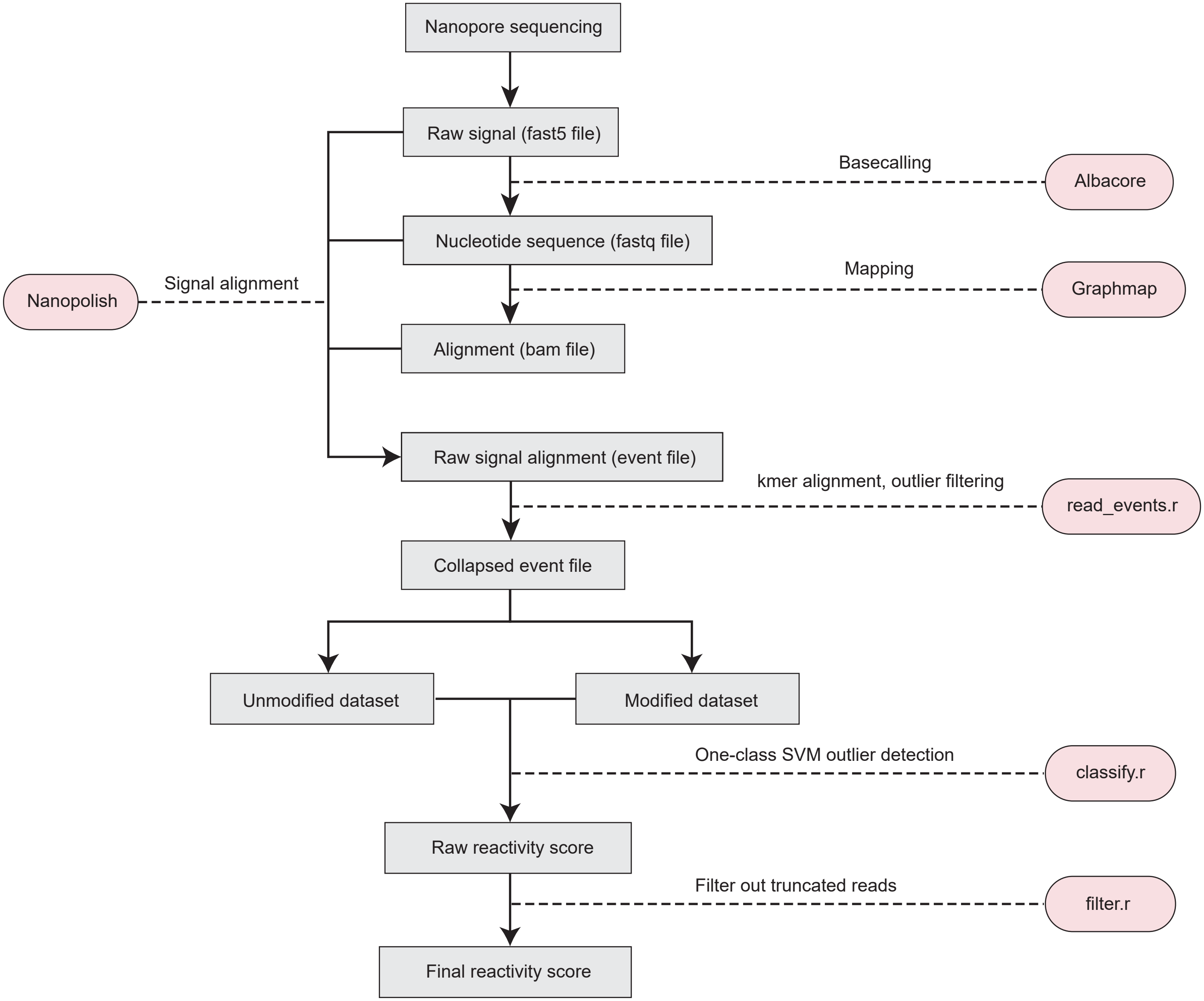

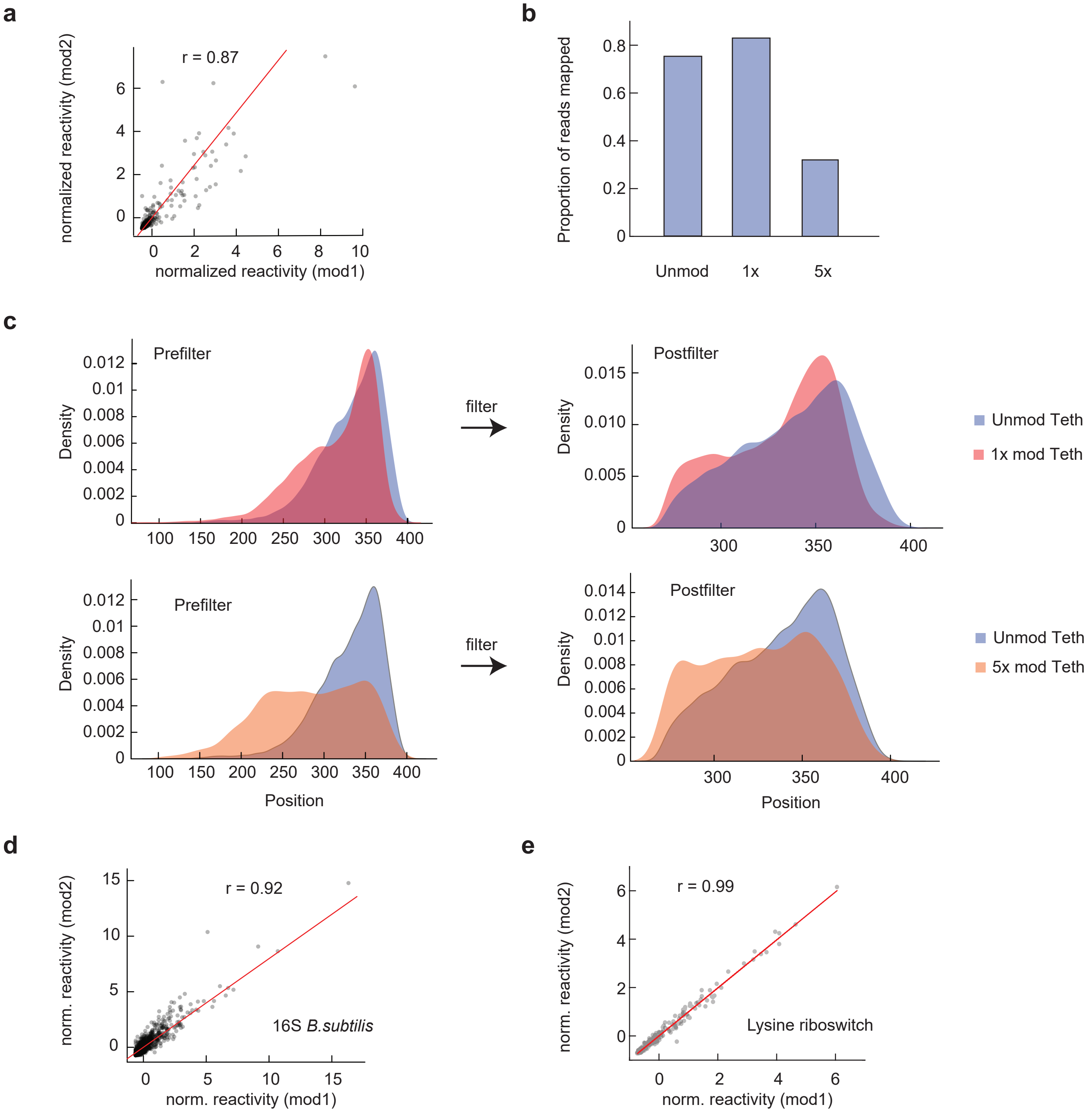

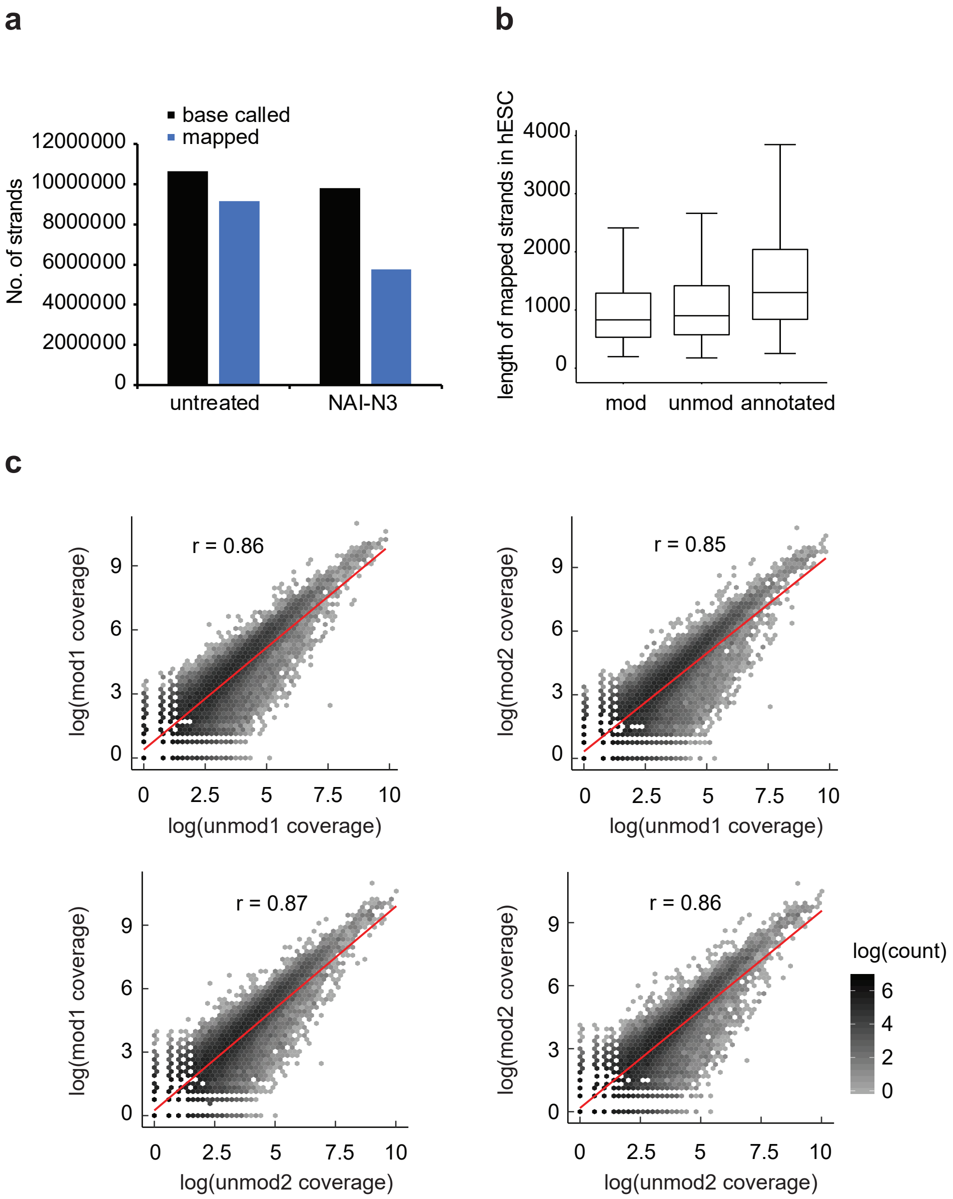

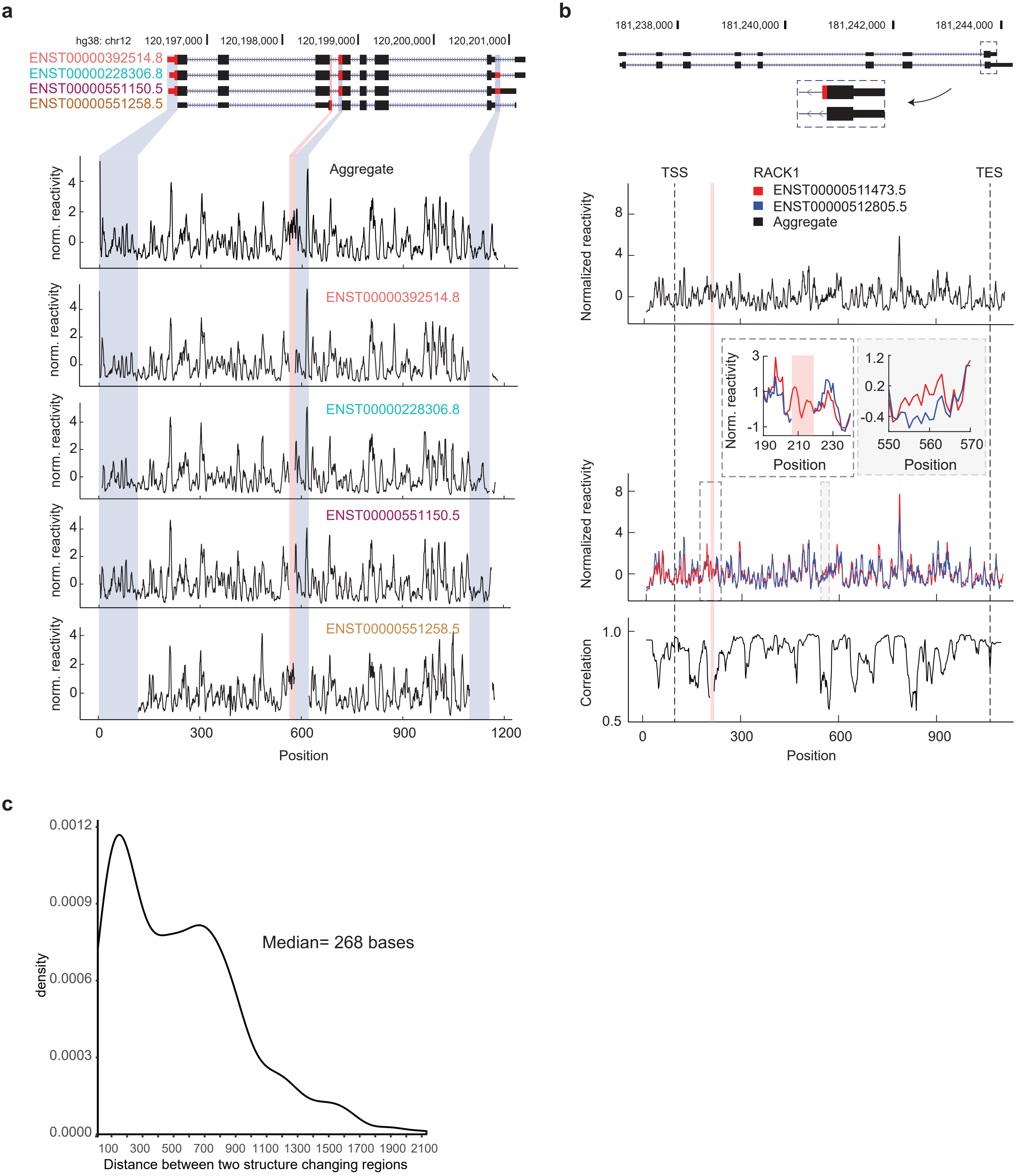

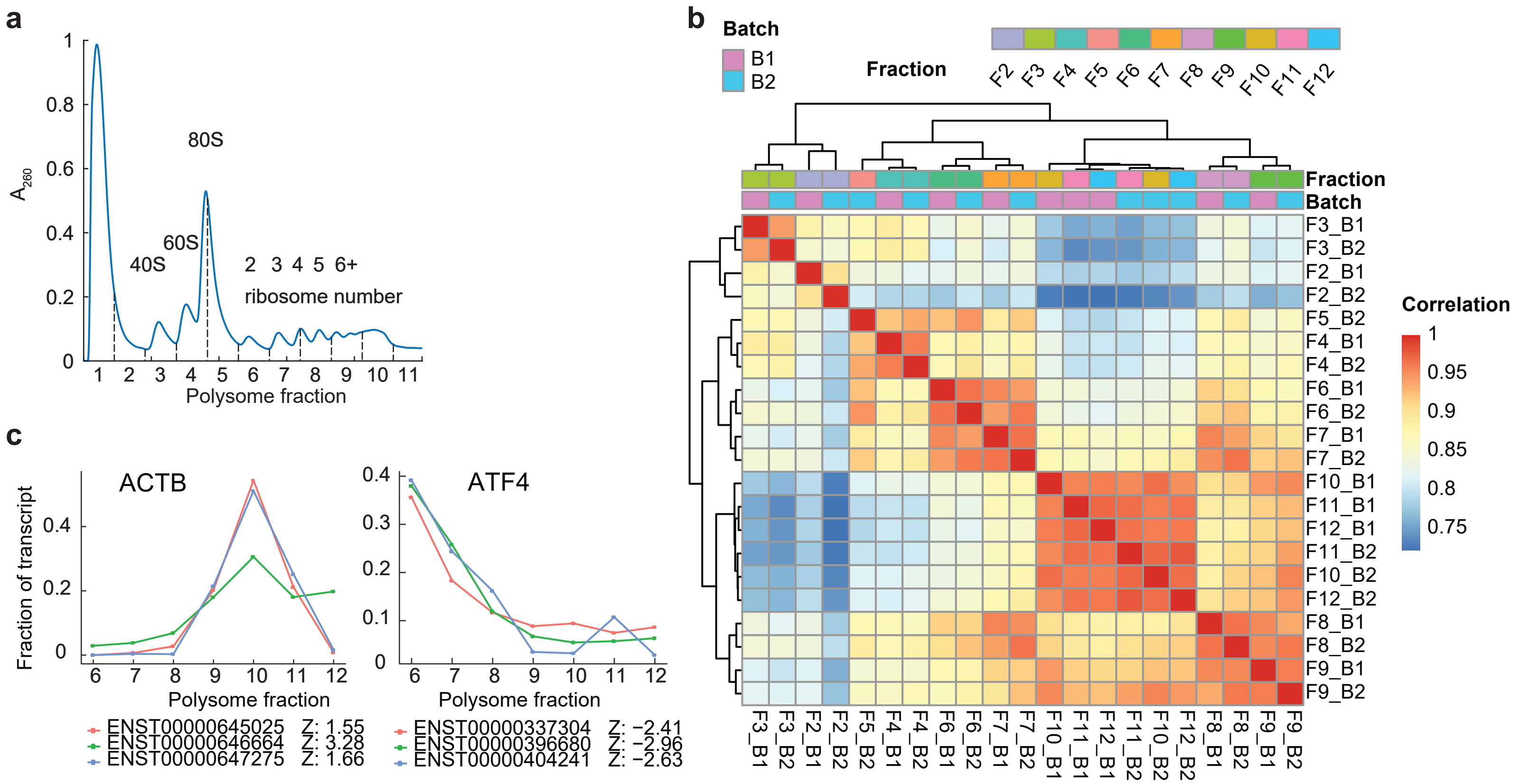
