## Supplemental table for "Direct RNA sequencing reveals structural differences between transcript isoforms"

Supplementary Table 1. Direct RNA sequencing statistics of individual RNAs

| Number | RNA | Condition | Basecalled | Mapped |
|---|---|---|---|---|
| 1 | Tetrahymena | Unmodified | 20149 | 15154 |
| 2 | Tetrahymena | NMIA | 36722 | 28427 |
| 3 | Tetrahymena | CMCT | 20149 | 15154 |
| 4 | Tetrahymena | DMS | 68463 | 13679 |
| 5 | Tetrahymena | NAI | 54289 | 6962 |
| 6 | Tetrahymena | NAIN3 | 51760 | 42107 |
| 7 | Tetrahymena | NAIN3 | 19606 | 16929 |
| 8 | 16S rRNA | Unmodified | 20678 | 13601 |
| 9 | 16S rRNA | 1x NAI-N3 | 52576 | 27173 |
| 10 | 16S rRNA | 5x NAI-N3 | 56985 | 15070 |
| 11 | 16S rRNA | 5x NAI-N3 | 55743 | 22787 |
| 12 | Lysine riboswitch | Unmodified | 49407 | 47202 |
| 13 | Lysine riboswitch | 1x NAI-N3 | 48454 | 39310 |
| 14 | Lysine riboswitch | 1x NAI-N3 | 64258 | 55706 |
| 15 | TPP riboswitch | Unmodified | 61413 | 10908 |
| 16 | TPP riboswitch | Unmodified | 5260 | 4185 |
| 17 | TPP riboswitch | Unmodified | 11607 | 1469 |
| 18 | TPP riboswitch | 1x NAI-N3 | 52608 | 6063 |
| 19 | TPP riboswitch | 1x NAI-N3 | 26039 | 4993 |
| 20 | TPP riboswitch | 1x NAI-N3 with 100uM TPP | 64023 | 9832 |
| 21 | TPP riboswitch | 1x NAI-N3 with 100uM TPP | 17098 | 1343 |
| 22 | TPP riboswitch | 1x NAI-N3 with 100uM TPP | 50967 | 8123 |
| 23 | TPP riboswitch | 1x NAI-N3 with 750nM TPP | 49313 | 44252 |
| 24 | TPP riboswitch | 1x NAI-N3 with 500nM TPP | 46748 | 42269 |
| 25 | TPP riboswitch | 1x NAI-N3 with 250nM TPP | 41442 | 36955 |

**Supplementary Table 2. Sequencing statistics for hESC H9 cells**

| No. | Flowcell ID | Condition | Cell line | Basecalled | Mapped |
|---|---|---|---|---|---|
| 1 | FAH15784 | Untreated | H9 | 487942 | 254527 |
| 2 | FAH45827 | Untreated | H9 | 35718 | 26023 |
| 3 | FAH64107 | Untreated | H9 | 486255 | 415446 |
| 4 | FAH93704 | Untreated | H9 | 1117255 | 986653 |
| 5 | FAJ05033 | Untreated | H9 | 1044162 | 905462 |
| 6 | FAJ05273 | Untreated | H9 | 911857 | 782933 |
| 7 | FAK14702 | Untreated | H9 | 766556 | 689157 |
| 8 | FAK14901 | Untreated | H9 | 1014604 | 899038 |
| 9 | FAK41492 | Untreated | H9 | 1121660 | 976053 |
| 10 | FAK41933 | Untreated | H9 | 268503 | 229139 |
| 11 | FAK41949 | Untreated | H9 | 843189 | 718016 |
| 12 | FAK43432 | Untreated | H9 | 1142411 | 1015028 |
| 13 | FAK45693 | Untreated | H9 | 688074 | 611199 |
| 14 | FAK46537 | Untreated | H9 | 571446 | 500465 |
| 15 | FAK48878 | Untreated | H9 | 157555 | 140522 |
| 16 | FAH45827 | NAI-N3 | H9 | 67911 | 44820 |
| 17 | FAH64127 | NAI-N3 | H9 | 361039 | 199610 |
| 18 | FAJ05272 | NAI-N3 | H9 | 694649 | 371316 |
| 19 | FAK10667 | NAI-N3 | H9 | 401243 | 227569 |
| 20 | FAK10669 | NAI-N3 | H9 | 706247 | 399015 |
| 21 | FAK10927 | NAI-N3 | H9 | 593035 | 331011 |
| 22 | FAK10971 | NAI-N3 | H9 | 176522 | 101259 |
| 23 | FAK11292 | NAI-N3 | H9 | 496013 | 270714 |
| 24 | FAK14654 | NAI-N3 | H9 | 215676 | 140456 |
| 25 | FAK14666 | NAI-N3 | H9 | 292093 | 161007 |
| 26 | FAK14675 | NAI-N3 | H9 | 874006 | 578772 |
| 27 | FAK15052 | NAI-N3 | H9 | 409843 | 216900 |
| 28 | FAK15331 | NAI-N3 | H9 | 223687 | 146720 |
| 29 | FAK24356 | NAI-N3 | H9 | 647902 | 362296 |
| 30 | FAK26020 | NAI-N3 | H9 | 530745 | 319918 |
| 31 | FAK27129 | NAI-N3 | H9 | 609219 | 367314 |
| 32 | FAK27172 | NAI-N3 | H9 | 851652 | 481390 |
| 33 | FAK27760 | NAI-N3 | H9 | 571745 | 351367 |
| 34 | FAK33302 | NAI-N3 | H9 | 455370 | 295181 |
| 35 | FAK38467 | NAI-N3 | H9 | 615171 | 385374 |

Supplementary Table 3. Primers for direct RNA sequencing and structure probing

| Primer name | Primer Sequence (5'->3') | Comments |
|---|---|---|
| Oligo A | /5PHOS/GGCTTCTTCTTGCTCTTAGGTAGTAGGTTC | General primer for annealing with Oligo B to form adapter replacing RTA |
| Oligo B TPP | GAGGCGAGCGGTCAATTTTCCTAAGAGCAAGAAGAAGCCACGGATGGAC | Oligo B for annealing to Oligo A |
| Oligo B Tetrahymena | GAGGCGAGCGGTCAATTTTCCTAAGAGCAAGAAGAAGCCCGAGTACTCC | Oligo B for annealing to Oligo A |
| Oligo B Lysine riboswitch | GAGGCGAGCGGTCAATTTTCCTAAGAGCAAGAAGAAGCCAGAAAGAGCG | Oligo B for annealing to Oligo A |
| Oligo B 16S | GAGGCGAGCGGTCAATTTTCCTAAGAGCAAGAAGAAGCCAGAAAGGAGG | Oligo B for annealing to Oligo A |
| Tetrahymena Kinase R99 | TCCCGCAATTTGACGGTCTTGC | Primer for kinasing for structure probing of Tetrahymena RNA |
| Tetrahymena Kinase R273 | CTTCCCCGACCGACATTTAG | Primer for kinasing for structure probing of Tetrahymena RNA |
| Lys Kinase R191 | AGAAAATGATTTCTTGACAGCC | Primer for kinasing for structure probing of Lysine riboswitch RNA |
| 16S Kinase R274 | ACGCATCGTCGCCTTGGTGA | Primer for kinasing for structure probing of 16S RNA |
| 16S Kinase R488 | CTTTCTGGTTAGGTACCGTC | Primer for kinasing for structure probing of 16S RNA |
